# Skill acquisition increases myelination and strengthens functional connectivity in the sensorimotor circuit of the adult rat

**DOI:** 10.1101/661546

**Authors:** Cassandra Sampaio-Baptista, Antoin de Weijer, Annette van der Toorn, Willem M. Otte, Anderson M. Winkler, Alberto Lazari, Piergiorgio Salvan, David M. Bannerman, Rick M. Dijkhuizen, Heidi Johansen-Berg

**Author notes:** Joint-first author (Cassandra Sampaio-Baptista, Antoin de Weijer).

## Abstract

The effects of skill acquisition on whole-brain structure and functional networks have been extensively investigated in humans but have yet to be explored in rodents. Forelimb reaching training in rodents results in well-established focal functional and structural reorganization within the motor cortex (M1) and cerebellum, indicating distributed alterations in both structure and function. However, it is unclear how local alterations in structure and function relate to distributed learning-related changes across motor networks. Here we trained adult rats in skilled reaching and used multimodal whole-brain *in vivo* MRI to assess both structural and functional plasticity over time.

We detected increases in a myelin-related MRI metric in white matter, cortical areas, and to a lesser extent in the cerebellum, paralleled by strengthened functional connectivity between M1 and cerebellum, possibly reflecting a decrease in cerebellum inhibition over M1. Skill learning therefore leads to myelin increases in pathways that connect sensorimotor regions, and in functional connectivity increases between areas involved in motor learning, all of which correlate with performance. These findings closely mirror previous reports of network-level changes following motor learning in humans and underlines the correspondence between human and rodent brain circuits for motor learning, despite important differences in the anatomy of physiology of movement circuits between species.

## INTRODUCTION

Motor skill learning induces local functional and structural changes in the rodent brain that can be detected with a variety of *ex vivo* and *in vivo* methods such as electrophysiology^1,2^, immunohistochemistry^2,3^, transcranial two-photon microscopy^4,5^ and calcium imaging^6,7^. For instance, skilled reaching training has been found to reorganize the representation of the trained paw and to induce synaptogenesis within the motor cortex in the late stages of learning^2,8^. Furthermore, spine formation in motor cortical regions has been detected in the early learning stages within hours and days^4^. In addition to such local changes within motor cortex, this task also results in changes in white matter (WM) structure underlying motor cortex, in part reflecting higher WM myelin after training^3^. This is consistent with the emerging notion that learning triggers myelin plasticity^9–11^ and suggests that effects of skilled reaching training extend beyond local motor cortical changes. Accordingly, cerebellar disruption impairs forelimb reaching movements^12,13^. Additionally, studies of complex motor learning in rodents on an obstacle course have shown that training results in alterations in the cerebellum such as astrocyte hypertrophy, increased complexity of the dendritic tree and increased number of synapses^14–17^. This is to be expected given that motor learning depends not only on motor cortical plasticity but also on functional interactions between distributed cortico-cerebellar and cortico-striatal circuits^18–21^. This has been demonstrated in human motor learning, where a combination of cortical, cerebellar and subcortical brain areas have been found to functionally interact at different learning stages and during different task components (for review see^22^). How such distributed learning-related changes across motor networks relate to local alterations in structure and function is unclear as local changes have been primarily studied in rodent models, while changes across motor circuits have commonly been explored in humans.

Non-invasive neuroimaging techniques, such as magnetic resonance imaging (MRI), can provide indirect assessment of changes in distributed functional interactions as they allow for simultaneous whole brain measurements in both humans and animals. For instance, resting-state fMRI (rs-fMRI) can be used to perform functional connectivity analysis, which refers to the statistical correlation between brain regions’ signal time-courses, to monitor how brain areas interact with each other, even when the subject is at rest and not performing a task inside the scanner. Short-term motor learning^23,24^ and long-term complex skill learning interventions^25^ modulate motor resting-state connectivity in humans. However, rs-fMRI functional connectivity changes with skill learning have not yet been assessed in rodents.

Skilled reaching training offers a powerful paradigm with which to explore distributed functional and structural changes with whole brain MR imaging, given that the existing literature provides extensive evidence for local structural and functional motor cortical changes^2,4,8^, functional modulation of thalamocortical projections^26^ and local cerebellar functional and structural reorganization^27^. This approach allows us to assess plastic changes within the same animals over time at the whole brain level, and to relate the findings to previous studies in both rodent and human. Additionally, applying whole-brain methods commonly used in humans to rodents, can inform future assessments with invasive approaches that are unavailable in human experiments, which can potentially clarify the underlying cellular mechanisms of human learning and behaviour.

We trained adult rats in a skilled reaching task and used multimodal *in vivo* MRI methods to assess both structural and functional changes. In a previous postmortem diffusion tensor imaging (DTI) study with the same motor learning paradigm, we found group differences in DTI metrics consistent with higher myelination along with higher myelin expression, as measured by immunohistochemistry^3^. Here, we first tested if we could detect increases in myelination non-invasively and *in vivo* by employing magnetization transfer ratio (MTR), a technique that can be used as an indirect assessment of myelin content^28,29^. Secondly, we monitored functional changes at the whole brain level with resting-state fMRI.

We specifically tested for functional connectivity alterations between the motor cortex and the rest of the brain.

## RESULTS

### Behavioural Results

Adult male Lister hooded rats (total = 36), (3 months old, 250 – 450 g) were randomly assigned to one of three experimental conditions: skilled reaching task (SRT), unskilled reaching task (URT) and a no training control condition (CC). Skilled reaching^30^ consisted in training rats to reach and grasp a sucrose pellet for 11 days. Rats assigned to the unskilled version of the task^2^ were trained to reach, but not to grasp as the pellet is put out of their reach, and were fed sucrose pellets on average every 5 reaches (see more details in Methods).

For the SRT group, a repeated-measures (RM) – ANOVA on reaching accuracy scores over 11 days of training showed a significant effect of day (F _(degrees-of-freedom: 2.247, 24.72)_ = 6.611; p < 0.01), confirming that accuracy improved over time as the rats learned the task (Fig. 1A). A subgroup of rats (n = 6) was tested 4 weeks after the last day of training. There were no significant differences between the 11^th^ day of training and this follow up test (t _(5)_ = −0.160, p = 0.879, 2-tail), showing that the rats performed at the same level 4 weeks after the training had ceased. At this same follow-up time-point, the URT group (n = 6) was also tested on the skilled reaching version of the task. Their performance was significantly worse than the SRT group performance on the same day (t _(10)_ = 2.853, p < 0.05, 2-tail), confirming that the skilled reaching training led to a clear and lasting improvement in reaching performance compared to the unskilled reaching group (Fig. 1A).

**Fig. 1.**
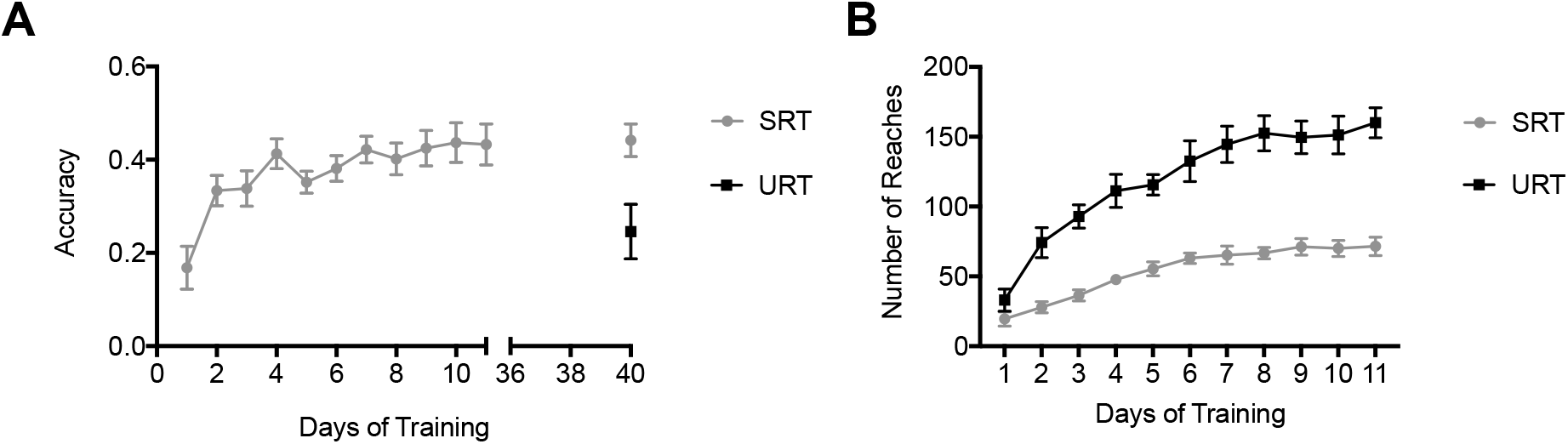
Behavioural Results. A Mean accuracy scores for SRT animals for all training days (grey) (n = 12). A subgroup of SRT (n = 6) and URT (black) (n = 6) rats were tested 4 weeks after the last training day on the skilled version of the task. B Mean number of reaches per day for SRT and URT (n = 12 animals per group). Error bars represent standard error.

Reaches are made during both the skilled and unskilled versions of the task. Mixed design ANOVA on total reaches per day in both URT and SRT groups, over 11 days of training, revealed a main effect of group (F_(1,22)_ = 52.107; p = 0.001; Fig. 1B) with the SRT group reaching less overall since as they become more successful they need to make fewer reaches to retrieve a pellet. Therefore, any brain changes that are found to be greater in the SRT compared to URT group should relate to learning, whereas any brain changes that are greater in the URT compared to the SRT group should relate to limb activity. A main effect of day (F _(4.482, 98.61)_ = 42.683; p < 0.001; Fig. 1B) and an interaction between day and group (F _(4.432, 98.61)_ = 6.023; p < 0.001; Fig. 1B) were also detected for number of reaches, as both groups of animals learn that reaching results in pellets, and so their number of reaches increases over time, but the SRT group becomes increasingly accurate on the task and fewer reaches are necessary to retrieve the pellets.

### Imaging Results

#### Increases In MTR In WM Tracts After Skilled Reaching Training

All rats were scanned 24 hours after the pre-training phase and 24 hours after the 11 days of training. A subgroup of animals (n=6 for each group, total n=18) was also scanned 24 hours after the 4 weeks follow up reaching test.

To assess effects of motor skill learning on WM microstructure we jointly analysed structural measures derived from in vivo MRI. We performed a non-parametric combination (NPC) for joint inference analysis, as implemented in Permutation Analysis of Linear Models (PALM) tool^31^, on the MTR maps and the 4 DTI measures (fractional anisotropy (FA), mean diffusivity (MD), radial diffusivity (RD) and axial diffusivity (AD)), using the Fisher’s combining function. NPC works by combining test statistics or p-values of separate (even if not independent) analyses into a single, joint statistic, the significance of which is assessed through synchronized permutations for each of the separate tests (see Methods for more details). We tested for a concordant direction of effects across all measures, while allowing the assessment of the significance maps for each measure separately, with correction for the multiplicity of tests.

We tested for baseline differences between groups and found no significant effects, either for the joint test or partial tests for any of the measures.

To specifically test for effects of skilled reaching training, we tested for an interaction effect between time and group across all voxels in the WM skeleton. The Fisher test did not reveal a significant interaction effect when considering all measures together. However, the partial tests revealed a significant interaction effect for MTR between group and scan (p < 0. 05, corrected) driven by an increase in MTR in the SRT group between scan 1 and scan 2 (Fig. 2 A, B). This increase was localized to the contralateral hemisphere of the used paw and comprised areas that overlapped with the significant cluster previously reported in a separate *ex vivo* study of the same task in different groups of rats^3^ (see also supplementary data Fig. 1, Fig. 2). In addition, the mean change in MTR within this cluster was found to positively correlate with mean accuracy in the skilled task across animals in the SRT group (Spearman r = 0.7455, p = 0.01, Fig. 2 C).

**Fig. 2.**
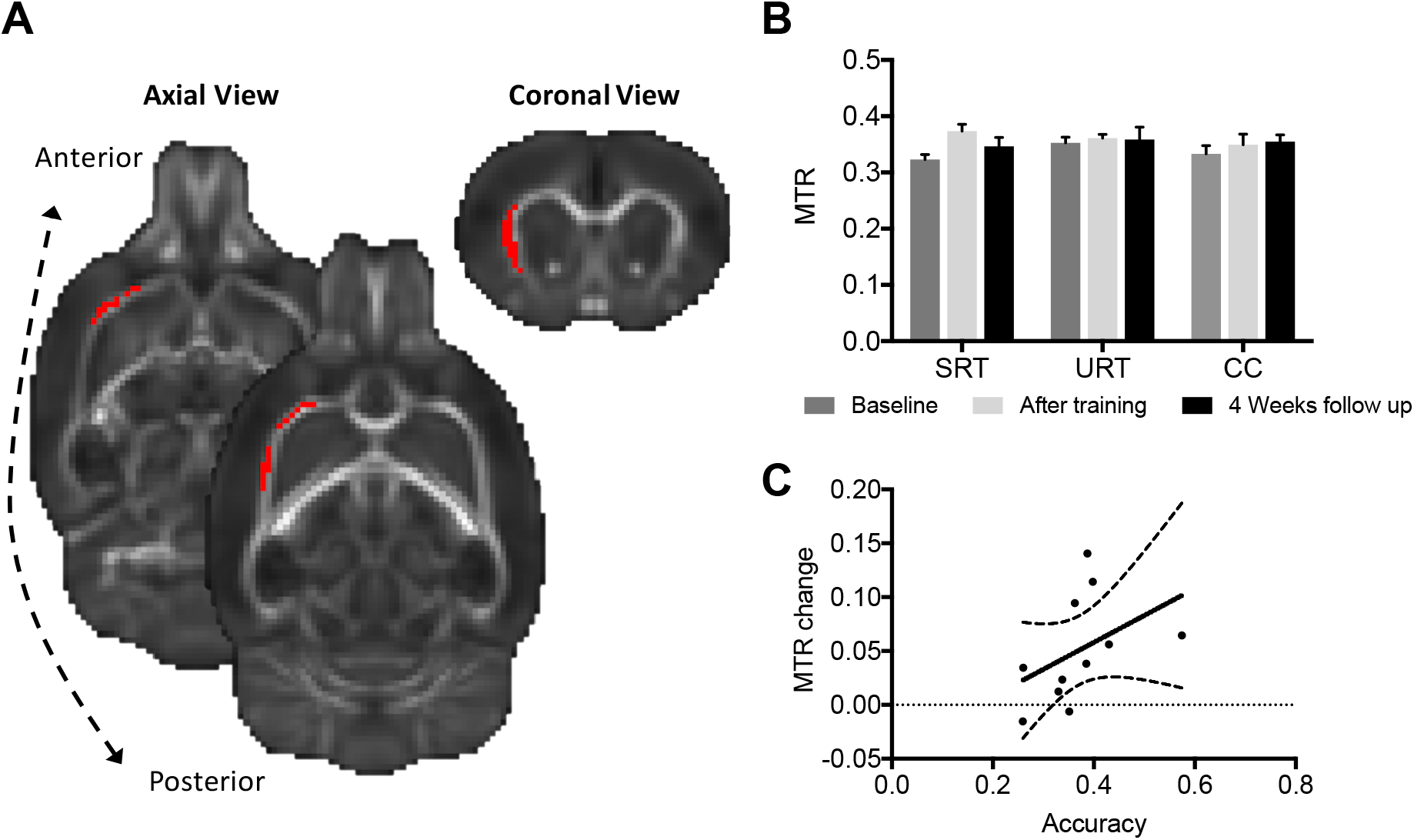
Effects Of Skilled Reaching Training On WM. ***A** Skilled reaching resulted in increases in MTR (red cluster) (p < 0.05, corrected). Significant clusters are superimposed on the mean FA template as this image has higher contrast than MTR and allows for better visualization. **B** Bar graph of mean MTR of the cluster illustrated in A per group* (Skilled reaching group (SRT), unskilled reaching group (URT) and caged control (CC)) *and time-point. This is shown to illustrate the direction of differences and not for statistical inference. Error bars represent standard error. **C** Scatter plot showing the correlation between mean accuracy in the SR task and mean change in MTR within the significant cluster shown in A (Spearman R = 0.7455, p = 0.01). Dashed lines represent standard error.*

#### Increases In MTR In Sensory Cortex After Skilled Reaching Training

Similarly, we analysed grey matter (GM) using MTR and MD and tested for joint effects across the 2 measures. MD can indicate changes in tissue density regardless of the structure orientation as, unlike WM, GM microstructure does not have dominant orientational structure at the spatial resolution employed here.

The Fisher test did not reveal a significant interaction effect when considering the two measures together. The partial tests revealed a significant interaction effect between time and group, driven by increases in MTR (p < 0.05, corrected) in the SRT group in GM regions (Fig. 3). The increases in MTR were seen primarily in S1, and S2 regions, and the striatum (Fig. 3). Just below the significance threshold a cluster in cerebellum (7-10 lobules including the paramedian lobule areas), was also identified (p = 0.07, corrected, Fig. 3A). No baseline differences between groups were identified.

**Fig. 3.**
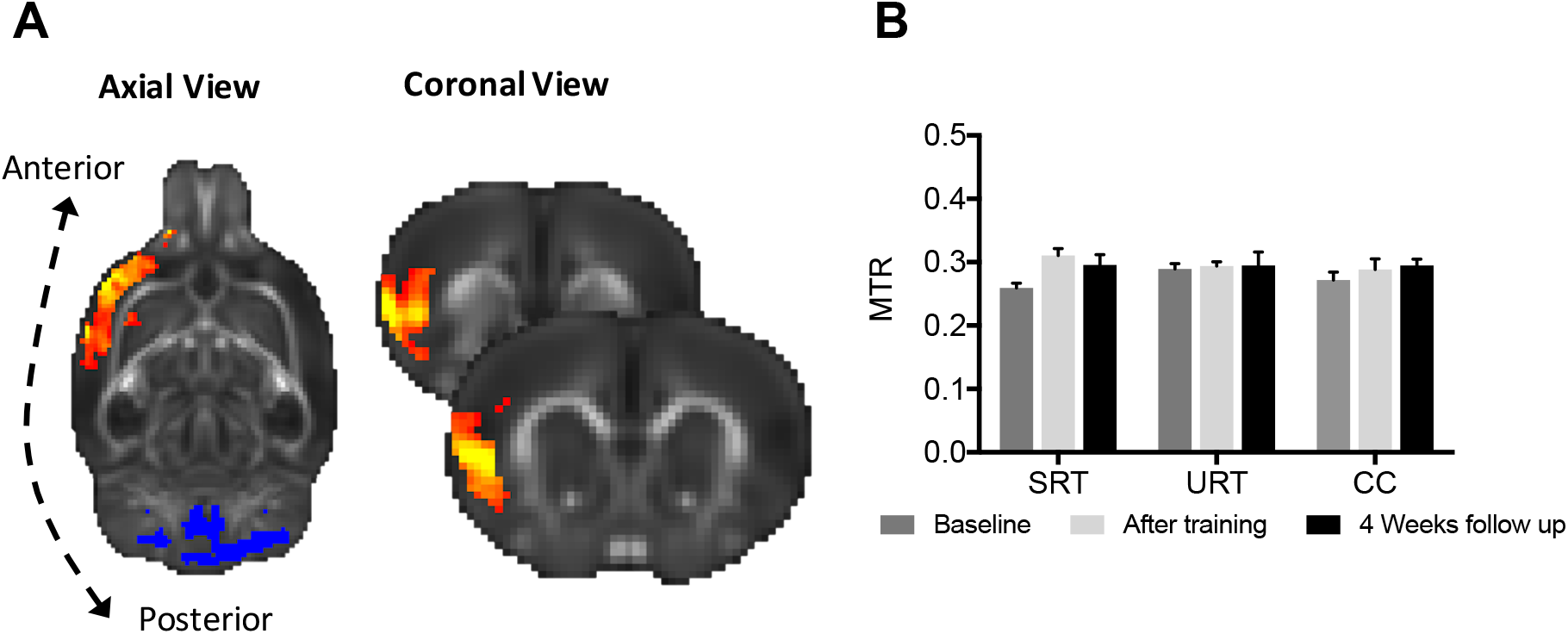
Effects of Skilled reaching training in GM. MTR (in yellow-red) was significantly increased in the SRT group (p < 0.05, corrected). A trend (p = 0.07, corrected) towards significant increases in MTR was also detected in the cerebellum (in Blue). Clusters are superimposed on the mean FA template as this image has higher contrast than MTR and allows for better visualization. **B** Bar graph of mean MTR of the significant cluster is shown to illustrate the direction of differences and not for statistical inference. Skilled reaching group – SRT; unskilled reaching group – URT; caged control – CC. Error bars represent standard error.

Similarly to the WM analysis, the mean change in MTR within the significant cluster in sensory areas was found to positively correlate with mean accuracy in the SR task for the SRT group (Spearman r = 0.7091, p < 0.05, not shown).

#### Structural Connectivity Results: WM Areas With Increased MTR Connect Functionally Relevant Areas For The Skilled Reaching Task

We used the baseline (n=36) diffusion data to perform tractography to identify the brain areas connected by the WM tract with significantly increased MTR (see Fig. 2). Widespread areas, including motor and sensory areas (Video 1), were found to be connected to the significant WM MTR cluster. The sensory areas identified by tractography partially overlapped with the GM regions (see Fig. 3) found to have significantly increased MTR after skilled reaching training.

**Video 1** *Red-yellow maps represent the mean tractography maps (n = 36). Dark blue areas represent S1/S2 and light blue areas represent M1/M2 (both derived from^32^).*

#### Functional connectivity: Increases in functional connectivity correlate with skilled reaching performance

We used seed-based voxel-wise analysis to assess functional connectivity changes between contralateral M1/M2 (derived from^32^ and represented in light blue in Fig. 5B) with the rest of the brain. M1/M2 was selected a *priori* as a seed region since it has been the main target of previous studies due to its pivotal role in the skilled reaching task^2,4,8,33^. No baseline differences were found between groups.

**Fig. 5.**
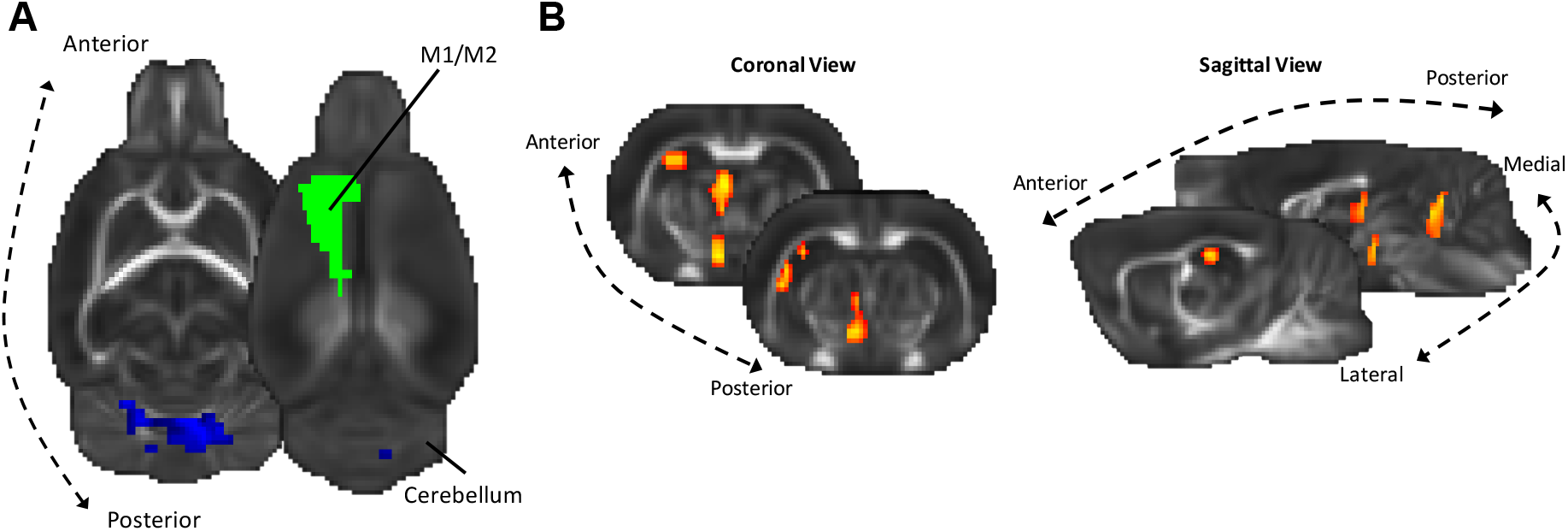
Seed-Based analysis results. **A** A seed region in M1M2 (green) was used to assess functional connectivity with the rest of the brain. A trend (p=0.09, corrected for multiple comparisons) towards increases in functional connectivity (represented by blue clusters) was found between the M1M2 seed and bilateral cerebellar areas in the SRT group compared to the URT group. **B** Within the SRT group, task performance correlated positively with connectivity changes (p < 0.05, corrected for multiple comparisons) between the M1M2 seed (shown in green in A), bilateral cerebellum and the hippocampus (represented by orange-red clusters). Significant clusters are superimposed on the mean FA template for visualization and anatomical comparison between modality results.

We tested for interaction effects between group and scan time-point and observed a trend towards an increase in functional connectivity in the SRT group compared to the URT group (p = 0.09, corrected) representing an increase in connectivity between M1M2 and bilateral anterior cerebellar areas in the SRT group (Fig. 5A).

In the SRT group, functional connectivity (fc) changes (Δfc – fc [after training] – fc [baseline]) between contralateral M1M2 and the rest of the brain were related to task performance (mean accuracy). Connectivity changes correlated positively (p < 0.05, corrected for multiple comparisons) between M1M2 and several brain areas such as bilateral anterior cerebellum, the hippocampus, as well as areas that appear to co-localise with thalamic and hypothalamic nuclei (Fig. 5B).

## DISCUSSION

Training on a skilled reaching task resulted in increases in MTR, a MRI-based metric that reflects myelin content, in white matter tracts that primarily connect sensory and motor regions, as well as in sensory cortical areas, and to a lesser extent in cerebellum. A trend for functional connectivity increases between motor areas and cerebellum was also seen after skilled reach training. The increase in MTR in both grey and white matter correlated with task performance, with greater increases in myelination being associated with higher performance. This is in accordance with our previous results showing a positive correlation between skilled reaching task performance and myelin expression, as measured by immunohistochemistry, with the same motor learning paradigm^3^. Our current results show that motor skill learning is associated with myelin-related structural changes in white matter pathways that connect task-relevant regions that also display increased intra-regional MTR and strengthened interregional functional connectivity.

Skilled reaching training resulted in greater strengthening of functional connectivity between motor cortex and cerebellum compared to the unskilled version of the task, and greater increases in cortico-cerebellar functional connectivity were associated with significantly better task performance. Earlier studies have provided evidence that the cerebellum, motor cortex and the caudate nucleus are important for forelimb reaching^12,13,34^. An increase in resting-state functional connectivity between M1 and cerebellum with motor learning has also been reported in humans. For instance, participants with greater motor sequence learning rate increases have stronger functional connectivity of the M1 and SMA with the cerebellum^35^. Increases in strength in the fronto-parietal network and the cerebellum network after 11 minutes of motor learning have also been detected^36^. This is further supported by EEG evidence showing increased resting-state connectivity within the cerebellum and between the cerebellum and motor cortex, after training in a force task^37^. While in baseline conditions, cerebellar cortex inhibits activity in motor cortex, via inhibition of excitatory outputs, motor learning in humans and animals reduces cerebellar inhibition over M1^27,38–41^. Hence, the increase in connectivity between motor and cerebellar areas detected here and in previous human studies, might reflect a decrease in inhibition of the cerebellum over M1, more specifically through disinhibition of the cerebellar nuclei that project to the thalamo-cortical circuit.

The importance of white matter plasticity and myelination to learning and plasticity during adulthood has been recently highlighted by a number of studies (for an overview see^11^). For instance, a recent study provided causal evidence that new myelin is required for novel motor acquisition using a conditional transgenic mouse model^10^. Using the same skilled reaching learning paradigm as used here, we previously found higher myelin basic protein (MBP) expression^3^ in pathways that co-localise with the MTR changes identified in the current study. Taken together these results provide the first evidence that MTR is sensitive to longitudinal changes in myelin in white matter. While previous studies have focused on the motor cortex, where a higher number of recently differentiated oligodendrocytes are found after motor learning^42,43^, here, using a whole-brain approach, we detected MTR increases in S1 and S2, and a trend in cerebellum, suggesting increases in myelination in these areas as well.

The MT effect measures macromolecule-bound proton exchanges with free water protons and has been used as an indirect assessment of myelin content given that most macromolecular content in the CNS is myelin. Validation studies have indicated that MT is sensitive to myelin in white matter lesions in multiple sclerosis^28^, but is not specific to myelin alone, as other changes can be due to edema^44^, inflammation and subsequent immune responses in these patients^45,46^. Given that in principle learning in healthy animals should not induce inflammation or edema, changes in MTR in the present study are highly likely to be related to myelination.

DTI differences were detected in the previous postmortem study, specifically higher FA in the SRT group^3^. In the current *in vivo* study, MTR proved to be a more sensitive measure than DTI. There are a few methodological differences that might explain the lack of sensitivity of the DTI measures in the current *in vivo* study. Our previous study included a larger sample, used *postmortem* imaging (allowing for longer scanning times and higher spatial resolution), and did not include MTR. DTI and MT rely on different MR effects and are effectively measuring different (though partially overlapping) tissue properties. While MTR is preferentially sensitive to myelin, DTI measures water diffusion across space and is sensitive to a number of tissue microstructure properties such as fibre orientation and density, axonal caliber, myelination and likely to other glial cells (for an overview see^47^). Nevertheless, the WM MTR increases identified in the current study partially overlap with the higher WM FA identified in the previous study^3^. Additionally, region of interest analysis of the previous study tract revealed significant increases in both FA and MTR in the SRT group only (although a group by time interaction was not significant, see supplementary data Fig. 1).

In summary, the current study found evidence for myelin increases in pathways connecting sensorimotor regions and for task performance-relevant increases in functional connectivity in association with skilled reaching training. These findings hint that myelination might occur in brain regions and in tracts that connect neurons engaged during the task. While in this study it is not possible to establish causal relationships between the detected structural and functional changes, future studies could assess if the learning-related synaptic changes occurring both in the cerebellum and motor cortex trigger myelination of the associated axons and in turn relate these different components to performance improvements such as speed or accuracy of the reach.

These findings in rodents closely mirror previous reports of network-level changes following motor learning in humans^25,36,48^. Detecting such network-level changes in association with learning in rodents underlines the correspondence between human and rodent brain circuits for motor learning, despite important anatomical differences in the physiology of movement circuits between species^49^. The whole-brain imaging approaches used in the current study highlight involvement of a distributed network of motor areas that could be further interrogated using invasive electrophysiological recordings, genetic manipulation or histological approaches unavailable in human experiments.

## MATERIAL AND METHODS

Experiments were approved by the committee for Animal Experiments of the University Medical Center Utrecht, The Netherlands (protocol number 2013.I.11.087) and were performed in accordance with the guidelines of the European Communities council directive. Adult male Lister hooded rats (total = 36), (3 months old, 250 – 450 g) (Harlan, Bicester, UK) were housed in groups of 3 (one per experimental condition) in standard laboratory conditions under a 12-h light/12-h dark cycle at 22-24°C temperature.

Before experiments all animals were given ad libitum access to typical food and water. A week before the start of the behavioural training all rats were food-deprived to 85% – 90% of their free-feeding weight (averaged over 3 consecutive days). Weight was closely monitored to ensure the animals did not fall below 85% of their original, free-feeding level. All animals were handled daily and provided with sucrose pellets (Dustless Precision Pellets®, 45 mg, Sugar, BioServ) in their home cage to habituate them to this new food type. All behavioural training and testing was performed during the light phase. Animals were randomly assigned to one of three experimental conditions: skilled reaching task (SRT), unskilled reaching task (URT) and control condition (CC) that consisted of caged controls.

### Behavioural training

The behavioural paradigm followed previously published guidelines on a single-pellet reaching task^2^ and are described in detail elsewhere^3^.

Prior to behavioural training all animals underwent pre-training sessions of 30 min/day for 3 days. A container filled with sucrose pellets (45 mg) (Bioserv, Frenchtown, NU, USA) was placed in front of the cage within easy reach. The pre-training stopped when the animal had retrieved 10 pellets or 30 minutes had passed.

After three days of pre-training, only the SRT and URT animals were further trained on the single-pellet-reaching task for 15 minutes daily for 11 consecutive days.

In the SRT training sessions a sugar pellet was placed within reach of the animal’s preferred limb (identified during the pre-training sessions). The animal had 5 attempts to successfully grasp the pellet, which together constituted one trial. Successful reaches were scored when an animal grasped the pellet and guided it to its mouth without dropping it.

To control for increased motor activity of the forelimbs, URT animals were placed in the same conditions as the SRT group but the pellet was placed out of reach, preventing the animals from developing reaching/grasping skills. To keep the animals motivated a pellet was dropped inside the cage periodically (every 5 attempts on average).

Four weeks after the end of training we tested both the SRT and URT groups in the skilled version of the task.

The CC animals did not receive any training after completion of the 3 days of pre-training but were handled daily and were weight-monitored and received a corresponding amount of sucrose pellets in their home cage.

### Behavioural measures and statistical analysis

The accuracy score was calculated as: number of successful retrievals / total number of reaches. Number of total reaches corresponds to the sum of all reaches per day. A repeated measures ANOVA (RM – ANOVA) was used to investigate improvement in accuracy scores over time in the SRT group. A separate Mixed design ANOVA was run to investigate differences in total number of reaches between SRT and URT groups. Sphericity was assessed and when appropriate Greenhouse-Geisser degrees of freedom were applied.

To confirm the absence of skill learning in the URT group, accuracy scores on the 4 weeks follow up period, when both groups were tested, were compared using a Student’s t-test. Mean accuracy was used to calculate behavioural correlations with imaging measures.

### MRI acquisition

MRI measurements were conducted on a 4.7 T horizontal bore MR system (Varian, Palo Alto, CA, USA) with a gradient coil insert (125 mm internal diameter; 500 mT/m maximum gradient strength with 110 μs rise time) (Magnex Scientific, Oxford, UK). A Helmholtz volume coil (90 mm diameter) and an inductively coupled surface coil (25 mm diameter) were used for signal excitation and detection, respectively. Rats were placed in an MR-compatible stereotactic holder and immobilized with earplugs and a tooth-holder.

Before MRI the animals were endotracheally intubated and mechanically ventilated with 2.0% isoflurane in a mixture of O2/air (1/2 volume/volume; 55 beats/min). During MRI, blood oxygen saturation and heart rate were continuously monitored by a pulse oximeter with the probe positioned on a hindpaw. In addition, expired CO2 was continuously monitored with a capnograph, and body temperature was maintained at 37.0 ± 0.5 C using a feedback-controlled heating pad.

Exactly 10 minutes prior to rs-fMRI acquisition, end-tidal isoflurane anesthesia concentration was reduced to, and maintained at 1.0%. At this level of isoflurane anesthesia, coherence of low-frequency BOLD signal fluctuations between functionally connected regions has been shown to be preserved^50^.

Resting-state fMRI was performed using a T2*-weighted single-shot gradient echo EPI sequence (TR/TE=700/20 ms; 75° flip angle; 40×34 matrix; 0.8 mm isotropic voxels; FOV 32 × 27.2 mm^2^ coronal, 17 slices; 850 BOLD images).

A diffusion-weighted four shot spin-echo EPI sequence was acquired (TR/TE=3000/23 ms, matrix size 80 × 80, resolution 0.4 mm isotropic, 8 averages per volume, 30 volumes with optimized diffusion encoding directions, 3 images with no diffusion weighting, 25 slices, field of view 32 × 32 mm^2^, b= 1334 s/mm^2^).

Magnetization transfer ratio images were acquired with a four shot spin-echo EPI sequence (TR/TE 3000/8ms, matrix size 80 × 80, resolution 0.4 mm isotropic, 25 slices, field of view 32 × 32 mm^2^, offset pulse 5000Hz).

### MR preprocessing and statistical analysis

We carried out analyses with the FSL package (www.fmrib.ox.ac.uk/fsl)^51^.

### DTI and MTR preprocessing

DTI data were analyzed with FMRIB’s Diffusion Toolbox (FDT)^51^ Fractional anisotropy (FA), mean diffusivity (MD), radial diffusivity (RD) and axial diffusivity (AD) were estimated from the original DTI data with DTIFIT.

Magnetization transfer ratio (MTR) was calculated by subtracting the image without offset pulse (mtr_off) from the image with offset pulse (mtr_on), and dividing this by the image without offset pulse (mtr_off) ((MTR_off – MTR_on) / MTR_off). Before further analysis the baseline scan of one rat was excluded due to a large artefact.

The trained limb was consistently aligned across animals. The FA images were first aligned with linear transformations to an existing rat brain template to generate the study specific template. All FA maps were aligned with linear and non-linear transformations to the study specific template and averaged to generate the mean FA image. MTR data was coregistered with the FA data. The FA registration transformations were applied to the remaining diffusion metrics (MD, RD and AD) and MTR images. To analyse grey matter only, MD and MTR images were masked with a grey matter mask derived from the mean FA.

We employed Tract-Based Spatial Statistics (TBSS)^52^ for the analysis of WM. The skeleton was extracted and thresholded at a conservative FA value of 0.36 to contain the main tracts while excluding spurious tracts^3^. Finally, the FA values of the tract centers were projected onto the skeleton for each rat brain and statistically analysed. MD, RD, AD and MTR values for the same voxels were also projected into the skeleton and voxel-wise analyses were performed.

### Tractography

We used tractography to identify the probabilistic connectivity map of the significant MTR cluster shown in Fig 2A. First, for each baseline scan (n=36), BEDPOSTX was used to automatically determine the number of estimated fiber populations per voxel and to fit estimates of principal diffusion direction for each population^53^. Then PROBTRACKX (5000 samples, 0.5 mm step length, 2000 steps, 0.2 curvature threshold) was used to follow these estimates in order to generate a probabilistic connectivity distribution, using the significant MTR cluster as a seed. The resulting individual maps were then overlapped across rats to produce the mean population probability map illustrating the connectivity of the MTR cluster.

### DTI and MTR Statistical analysis

We performed non-parametric combination (NPC), as implemented in the Permutation Analysis of Linear Models (PALM) tool^31^, for multimodal analysis. PALM is a tool that allows inference over multiple modalities, including non-imaging data, using non-parametric permutation methods, similarly to the *randomise* tool in FSL^54^, although offering a number of features not available in other analysis software, such as the ability for joint inference over multiple modalities, or multiple contrasts, or both together, while correcting FWER or FDR across modalities and contrasts^31^. NPC works by combining test statistics or p-values of separate analyses into a single, joint statistic, the significance of which is assessed through synchronized permutations for each of the separate tests, which implicitly accommodate any eventual lack of independence among them. Such a joint analysis can be interpreted as a more powerful, permutation-based version of the classical multivariate analysis of covariance (MANCOVA); differently than MANCOVA, however, NPC allows investigation of the direction of joint effects.

In WM, we assessed the joint and individual contribution of the 4 DTI measures (FA, MD, RD, and AD) and MTR while simultaneously correcting across the tests. For the WM skeleton analysis we used NPC with Fisher’s combining function, testing for joint effects across the 5 measures. A cluster-forming threshold of t > 2 and 5000 permutations were used to determine p-values family wise error rate (FWER) corrected for multiple comparisons (across all voxels and the 5 measures). The chosen cluster-forming t threshold was based on the degrees of freedom of the sample. Clusters with a corrected significance of p < 0.05 were deemed significant. For the GM analysis we focused on MD, which can indicate changes in tissue density regardless of the structure orientation, and MTR only. Similarly, we analysed GM using MD and MTR and tested for joint effects across the 2 measures. A cluster-forming threshold of t > 2 and 5000 permutations were used to determine p-values FWER-corrected for multiple comparisons (across all voxels and the 2 measures).

We tested for interaction effects between groups (SRT, URT and CC) and time-points (baseline versus after training). We calculated the differences between after training scans and baseline scans for each rat and used this difference image to directly test for interaction effects between groups. We also tested for baseline differences between groups.

A supplementary ROI analysis was done by extracting the mean FA and mean MTR values within the significant cluster found in^3^. We tested for group differences with a mixed design ANOVA, with group (SRT, URT and CC) as between subjects’ factor, and training day (Baseline vs After Training) as a within subject factor. Paired t-tests were performed to examine within group effects.

### Resting-state preprocessing and seed-based analysis

Preprocessing steps of the resting-state fMRI scans included brain extraction, removal of the first 50 volumes to reach a steady state, and motion-correction with MCFLIRT. At the preprocessing stage time-point two of one rat was excluded due to poor quality data. Motion-correction parameters were used as regressors for the resting-state signal (no linear detrending and global mean regression were performed).

After preprocessing, we performed seed-based correlation analysis using a mask of the contralateral M1M2 derived from a custom-built 3D model^55^ of the 5th edition of the Paxinos and Watson rat brain atlas^32^. We used the FSL tool “dual regression” to extract the time series and to regress those time-courses into the same 4D dataset to get a subject-specific set of spatial maps for each animal and time-point. The spatial maps were then used to perform voxel-wise analysis with non-parametric permutation testing as implemented by the FSL tool randomise, with a cluster-forming threshold of t > 2, and 5000 permutations were used to determine corrected p-values.

We first tested for baseline differences between groups (SRT, URT and CC). We then tested for interaction effects between groups and time-points (baseline versus after training). To do this, we calculated the differences between baseline and after training scans (after training – baseline) for each rat and used this difference image to directly test for interaction effects between groups and for effects of performance within the SRT group.

## Supporting information

Supplementary Results

## ACKNOWLEDGEMENTS

The authors would like to thank Ellen Thomas for her valuable help on the behavioural scoring. This work was supported by the Wellcome (WT090955AIA, 109062/Z/15/Z to AL and WT110027/Z/15/Z to H J-B) and the Netherlands Organization for Scientific Research (NWO-VICI 016.130.662 to RMD). The Wellcome Centre for Integrative Neuroimaging is supported by core funding from the Wellcome Trust (203139/Z/16/Z).

## CONFLICT OF INTEREST

We report no conflict of interest.

