## Supplementary Results for "Skill acquisition increases myelination and strengthens functional connectivity in the sensorimotor circuit of the adult rat"

*Region of interest analysis (ROI) analysis*

We performed a ROI analysis by extracting the mean values of FA and MTR within the cluster found significant in (Sampaio-Baptista et al., 2013). We tested for group differences with a mixed design ANOVA, with group (SRT, URT and CC) as between subjects' factor, and training day (Baseline vs After Training) as a within subject factor.

This ROI analysis of the mean FA revealed a main effect of day ($F_{(1,33)} = 9.086$; $p < 0.05$) but not a main effect of group or an interaction effect. As the main purpose of this analysis was to explore the extent to which the current results replicate previous findings, exploratory follow-up t-tests were performed despite the lack of interaction. Paired T-Tests within group showed a significant effect for the SRT group ($t_{(11)} = 3.985$, $p < 0.01$, 2-tail), but not for the URT ($t_{(11)} = 1.864$, $p = 0.089$, 2-tail) or CC ($t_{(11)} = 0.702$, $p = 0.497$, 2-tail) groups.

ROI analysis of the mean MTR revealed a main effect of day ($F_{(1,32)} = 4.578$; $p < 0.05$) but not a main effect of group or an interaction effect. Paired t-tests within group revealed a significant effect for the SRT group ($t_{(10)} = 2.501$, $p < 0.05$, 2-tail), but not for the URT ($t_{(11)} = 1.511$, $p = 0.159$, 2-tail) or CC ($t_{(11)} = 0.460$, $p = 0.654$, 2-tail) groups.

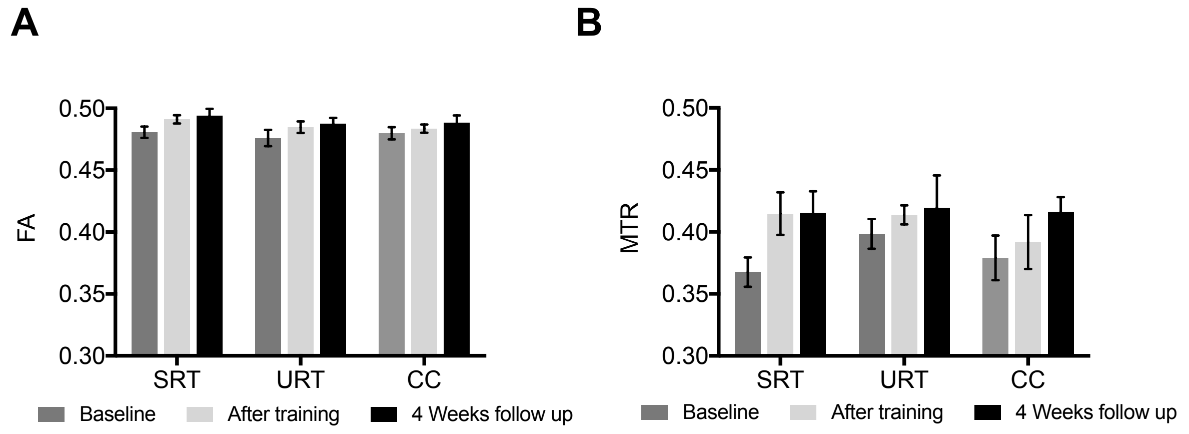


**Supplementary Fig. 1 ROI analysis A** A significant effect of day but not group or interaction was found for the FA. Paired t-tests revealed a significant effect for the SRT group but not for the control groups. **B** Mixed design ANOVA of the mean MTR revealed significant effect of day but not group or interaction. Paired t-tests revealed a significant effect for the SRT group but not for URT or SRT.

*Correlations between baseline MRI measures*

Baseline MTR is significantly anticorrelated with baseline RD (Shapiro-Wilk normality p=0.003; Spearman r=-0.434; p=0.0046, 1-tail), the higher MTR the lower RD. RD measures water restriction across the main axis which some authors interpret as being related to myelin and membrane integrity. As MTR is considered an indirect measure of myelin the correlation is in the expected direction. There was a trend towards a statistically significant positive correlation between baseline MTR and baseline FA (Spearman r=0.25; p=0.065, 1-tail), with the association between these two metrics being in the expected direction.
